# The infection dynamics of vertically transmitted viruses in a two-population model

**DOI:** 10.1101/284380

**Authors:** Jakob Friedrich Strauß, Marina Chauvet, Arndt Telschow

**Affiliations:** Institute for Evolution and Biodiversity, Westfalian Wilhelms-University, Münster, Germany, D-48149; University of Osnabrück, Osnabrück, Germany, D-49074

**Keywords:** host-parasite interaction, theory, vertical transmission, migration, mathematical modelling, bistability, *Drosophila*, sigma virus, geographic mosaic

## Abstract

There is a growing interest in vertically transmitted viruses. Field studies show that virus infection frequency often varies in space and time. In order to understand the spatial variation of virus prevalence, we designed a mathematical model describing the infection dynamics of a biparentally inherited virus in a two-population model. Model parameters are paternal and maternal transmission rates, cost of infection, and up to two migration rates. We investigated three different population structures: (1) a single panmictic host population, (2) two populations connected by unidirectional migration (mainland-island model), and (3) two populations connected by two-way migration. According to the results, the parameter space can be divided in three zones: (1) viral spread, (2) no viral spread, (3) bistability (in a single population) or stable infection polymorphism (in the two-population models). The finding of bistability is interesting and new. It shows that infected and uninfected populations can stably coexist, if migration is below a critical value, and if the cost of infection is moderately high. In summary, our results show that spatial structure of host populations is an important factor in determining geographical differences of vertically transmitted viruses.

## 2 Introduction

The transmission mode of a parasite is a key factor in determining its epidemiology and evolution (Ewald, 1987, 1994; Yamamura, 1993; Frank, 1996; Ebert et al., 1998, 2008). The two main modes are horizontal transmission (i.e., from individual to individual) and vertical transmission (i.e., from parent to offspring). There is good empirical evidence that vertically transmitted parasites are common in nature (Mims 1981; Duron 2008). Examples include seed-borne infections in plants (Mink, 1993; Maule and Wang, 1996), viral infections in mammals (Mims, 1981), and reproductive parasites in arthropods (Werren et al., 2008). The wide distribution of vertically transmitted parasites makes their study an important topic in ecology and evolution. Here, we focus on the infection dynamics of biparentally inherited parasites, with a special emphasize on spatial effects.

A case in point is sigma virus, a well-studied rhabdovirus, which is known to infect several *Drosophila* species (L’Heritier, 1958, 1971; Carpenter et al., 2007; Longdon et al., 2011a, 2011b). Its transmission is vertical through both sperm and egg, but usually paternal transmission is less efficient than maternal transmission (Longdon et al., 2011b; Wayne et al., 2011). Sigma virus reduces host fitness by lowering both fertility (Yampolsky et al., 1999) and survival in winter (Fleuriet, 1981). In addition, it causes to its host a CO2 sensitivity (Williamson, 1961). Field screenings have revealed that though the virus has spread worldwide, it infects only a low proportion of flies in nature (Williamson, 1961; Fleuriet, 1992; Yampolsky et al., 1999; Carpenter et al., 2007). This pattern was suggested to be the result of spatial structure and migration between subpopulations (Fleuriet 1981; Yampolsky et al., 1999).

There is substantial theoretical literature on vertically transmitted parasites. Fine (1975) derived a general model for the infection dynamics in a panmictic population, and found conditions under which a biparentally inherited virus can spread and persist in a host population. Later studies investigated the combined action of vertical and horizontal transmission (Busenberg and Cooke, 1988; Lipsitch et al., 1995), or connected vertical transmission to the evolution of mutualism (Yamamura, 1993). More recent work has focused on the particular host-parasite system of sigma virus and Drosophila (Yampolsky et al., 1999; Longdon et al., 2011; Wayne et al., 2011). However, an explicit spatial model of the infection dynamics of vertically transmitted parasites has not been analysed yet.

In the present study, we investigated the infection dynamics of a biparentally inherited virus. First, we re-analysed a single population model proposed by Longdon et al. (2011b), and found a zone of bistability, which was not described previously. We then extended this model to incorporate spatial structure and migration. For three different scenarios we performed spread simulations, fixpoint analyses, and parameter screenings in order to determine conditions for viral spread and persistence. Interestingly, we found a zone of a stable infection polymorphism, where neighbouring populations differ in infection frequency despite migration between them.

## 3 Model and Results

### 3.1 Model parameters and population structures

We describe the infection dynamics of a vertically transmitted virus in a discrete time step model with non-overlapping generations. We follow Flor et al. (2007) and compare four different population structures (see Fig. 1): (a) a single panmictic population; (b) a mainland-island model with an uninfected mainland; (c) a mainland-island model with an infected mainland; (d) a two population model with bidirectional migration. In cases (b)-(d), migration is described by a migration rate, m, which is defined as the fraction of the target population, which is replaced by individuals from the source population at each generation. The migration rate may range from 0 to 1. In the model with two-way migration, migration rates may differ in both directions. Then *m*_*X*_ corresponds to the fraction of population *X* that is replaced by migrants.

**Figure 1:**
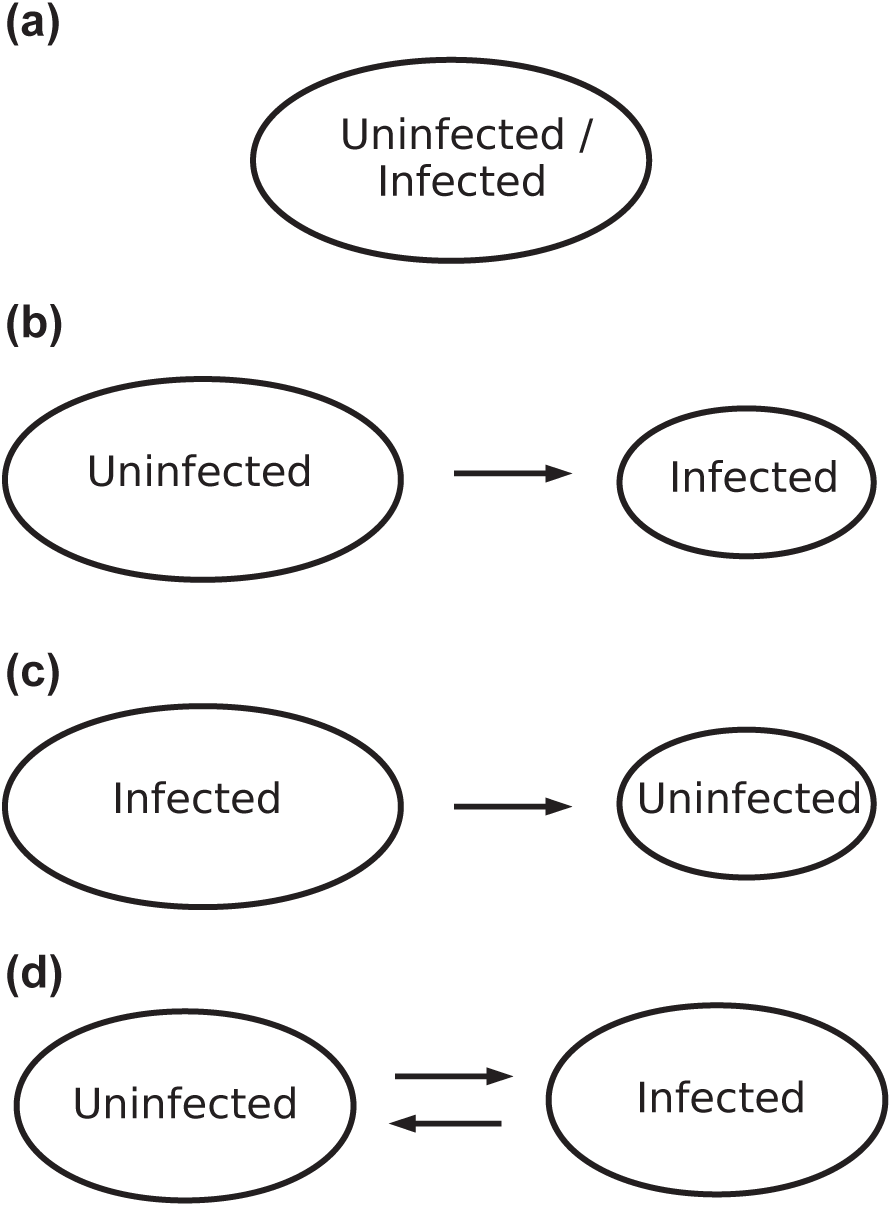
Model population structures and initial conditions. We analysed the infection dynamics of vertically transmitted viruses in four different scenarios: (a) a single panmictic host population with varying initial conditions for viral infection; (b) a mainland-island model with an uninfected mainland, and viral infection at equilibrium on the island as initial condition; (c) a mainland-island model with viral infection at equilibrium on the mainland, and no viral infection on the island as initial condition; (d) a two-population model with bidirectional migration, one population being uninfected and the other with viral infection at equilibrium as initial conditions.

Following Yampolsky et al. (1999) and Longdon et al. (2011b), we describe the viral infection dynamics in each population by three parameters: maternal transmission rate, *t*_*m*_, paternal transmission rate, *t*_*p*_, and relative fitness of infected hosts, *c*. All three parameters may take values between 0 and 1, and do not differ between the populations. The virus is assumed to be transmitted via both egg and sperm, but maternal and paternal transmission rates may differ. We define the maternal transmission rate *t*_*m*_ as the fraction of infected eggs among all eggs of an infected female, and the paternal transmission rate *t*_*p*_ as the fraction of infected sperm among all sperm of infected males. Viral infection is assumed to reduce both male and female fitness. The parameter *c* is defined as the relative fitness of infected individuals in comparison to uninfected individuals.

For solving equations and performing spread simulations, we used the software Mathematica (version 8.0.2.0, Copyright 1988-2011 Wolfram Research, Inc.) and R (version 2.14.1, 22.12.2011, Copyright 2011 The R Foundation for Statistical Computing).

### 3.2 Single host population

Let *P*_*i*_ and *P*_*i*+1_ denote the virus frequency in subsequent generations. Following Longdon et al. (2011b), the infection dynamics of vertically transmitted viruses in a single host population can be described by the following difference equation:

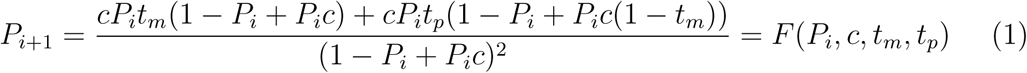

Equation 1 can be derived as follows (for a graphical illustration, see Appendix 6.1, Fig. 7). The numerator corresponds to the number of infected offspring, whereas the denominator corresponds to the total number of offspring. The first part of the numerator refers to the number of offspring, which received the infection from their mother, *cP*_*i*_*t*_*m*_(1 *-P*_*i*_ + *cP*_*i*_). Here, *cP*_*i*_ refers to the total number of eggs produced by infected females, *t*_*m*_ to the probability with which an egg is infected, and (1 *-P*_*i*_ +*cP*_*i*_) to the total amount of sperm from uninfected (1 *-P*_*i*_) and infected males (*cP*_*i*_). The second part of the numerator corresponds to the number of infected offspring, which have received their infection by their father and not by their mother, *cP*_*i*_*t*_*p*_(1 *-P*_*i*_ + *cP*_*i*_(1 *- t*_*m*_)). Here, *cP*_*i*_ refers to the number of infected males, *t*_*p*_ to the probability of paternal transmission, and (1 *-P*_*i*_ + *cP*_*i*_(1 *-t*_*m*_)) to the number of uninfected eggs produced by uninfected (1 *- Pi*) and infected females (*cP*_*i*_(1 *- t*_*m*_)). Finally, the denominator (1 *- P*_*i*_ + *cP*_*i*_)^2^ is the total number of progeny produced at generation*i* + 1, corresponding to the product of eggs coming from infected or uninfected parents (1 *- P*_*i*_ + *cP*_*i*_) and sperm coming from infected or uninfected parents (1 *- P*_*i*_ + *cP*_*i*_).

In a first step of the analysis, we investigated the temporal dynamics of the infection. Three qualitatively different outcomes are possible. With no or moderate costs of infection (0.67 < *c* ≤ 1), the virus spreads from low initial conditions to its equilibrium frequency, whereas for high costs of infection (*c* ≤ 0.67), the virus cannot invade the population from low initial frequency (Fig. 2a). Interestingly, we also found a zone of bistability, which was not described by previous authors. For example, with the same parameters as in figure 2a and *c* = 0.63, the virus cannot spread from low initial infection frequencies (in concordance with Fig. 2a), but can persist if it has already spread in the population (Fig. 2b).

**Figure 2:**
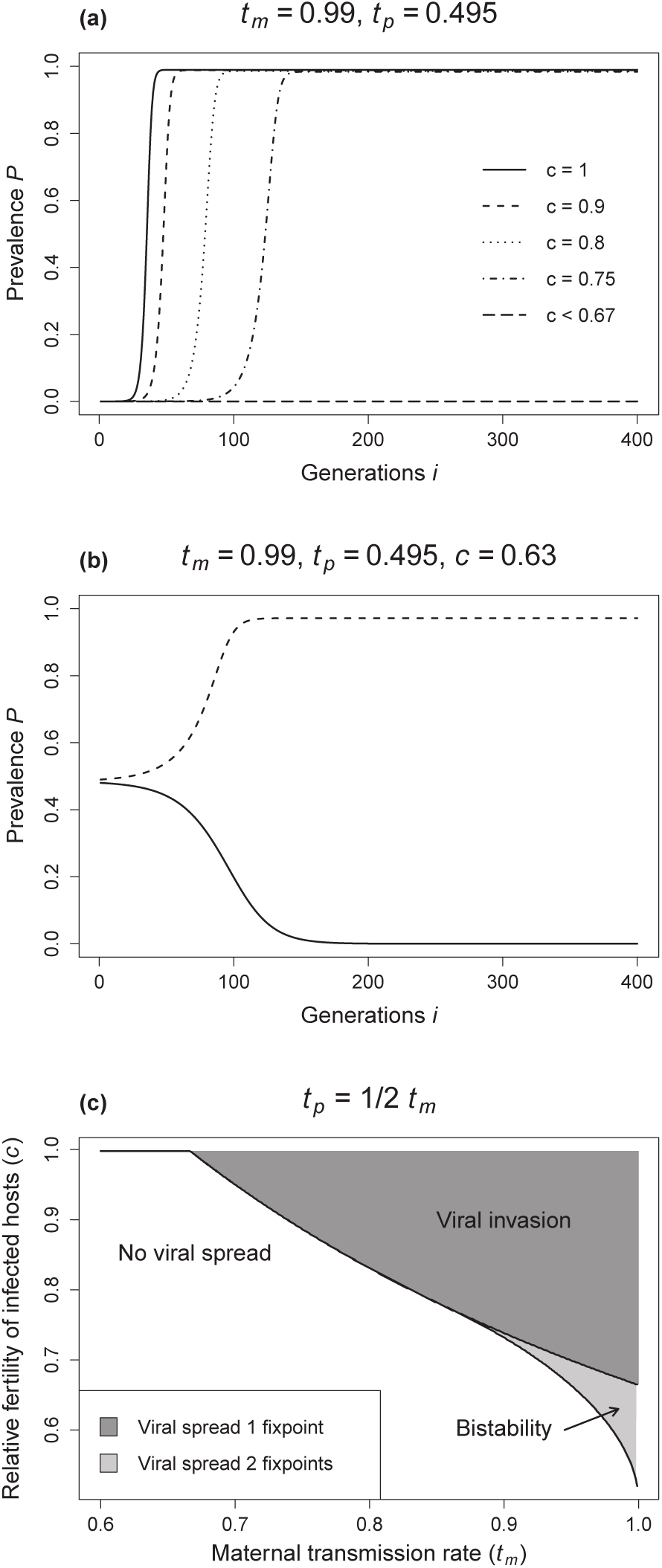
Infection dynamics in a single host population. (a) Viral spread is simulated by numerical iteration of equation 1, and an initial infection prevalence of *P*_0_ = 10^*-*6^. Transmission rates are *t*_*m*_ = 0.99 and *t*_*p*_ = 0.495, and the relative fitness *c* ranges between 0.67 and (b) Bistability occurs for *t*_*m*_ = 0.99, *t*_*p*_ = 0.495, and *c* = 0.63. Initial frequencies of viral infection of 0.48 or below result in viral extinction, whereas initial frequencies of 0.49 or above lead to viral spread and persistence. (c) Parameter screen of the three different zones: viral spread, viral extinction, and bistability.

In order to get a deeper understanding of the possible virus equilibrium frequencies, we performed a standard fixpoint analysis. Fixpoints were denoted by *P*^*^ and calculated by solving equation 1 for *P*_*i*+1_+1 = *P*_*i*_ = *P*^*^. This approach yields three fixpoints. The first, 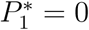, represents the state of no infection. It can be stable or unstable depending on the parameters. The other two fixpoints, 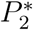 and 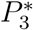, can be calculated analytically (see Appendix 6.1.1). There are three qualitatively different outcomes. First, if both 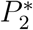 and 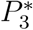 are neither not positive or purely complex, then 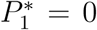 is the only stable fixpoint, representing the state where the infection can neither spread nor persist. Second, if 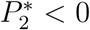 and 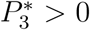, then 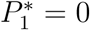 is unstable, and the stable fixpoint 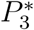 represents the equilibrium frequency of the viral infection. Third, if 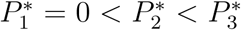, then bistability occurs with 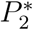 unstable, and 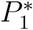 and 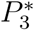 stable. Other circumstances cannot occur.

According to the fixpoint analysis, the parameter space can be divided in three zones: (1) no viral spread, (2) viral spread, (3) bistability. To illustrate the results, we performed a parameter screening for the scenario where paternal transmission rate is half of maternal transmission rate (Fig. 2c). As evident from the graph, the area of no viral spread is larger than the area of viral invasion, and the bistability area is smallest. The important insight from the analysis of the single population model is the existence of the bistability zone. This will be crucial for understanding the results in the next sections.

### 3.3 Mainland-island models

In this section, the infection dynamics is considered in a mainland-island model, where migration is unidirectional from the mainland to the island population. We analysed two cases, which differ with respect to their initial conditions. In the first, the mainland is uninfected and the infection is at equilibrium on the island. In the second, the infection is at equilibrium in the mainland, while the island is uninfected. For both scenarios, viral infection dynamics were analysed on the island, and it was investigated under which conditions differences in viral prevalence can stably persist in the face of migration. Following Flor et al. (2007), we call this situation stable infection polymorphism, and use the term critical migration rate (*m*_*c*_) for the highest migration rate, below which a stable infection polymorphism exists.

#### Uninfected mainland

Employing the function defined in equation 1, the prevalence of vertically transmitted viruses in the island population can be described by the following difference equation:

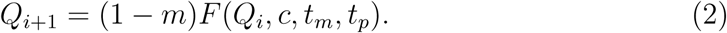

Here, *Q*_*i*_ and *Q*_*i*+1_ denote the virus frequencies in the island population in subsequent generations. As in the single population model, equation 2 has three fixpoints: 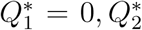 and 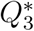 (for analytical solutions, see Appendix 6.1.1). These fixpoints are plotted against the migration rate *m* in figure 3a. Note that the fixpoints are only shown if they are real numbers. As can be seen, there is a critical migration rate, below which three real fixpoints occur, but above which 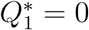 is the only real fixpoint. It can be shown that 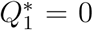 is always stable, and that 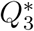 is stable if it is a real number. 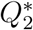 is in that case unstable and can be interpreted as a threshold frequency that determines whether an infection spreads or goes to extinction.

**Figure 3:**
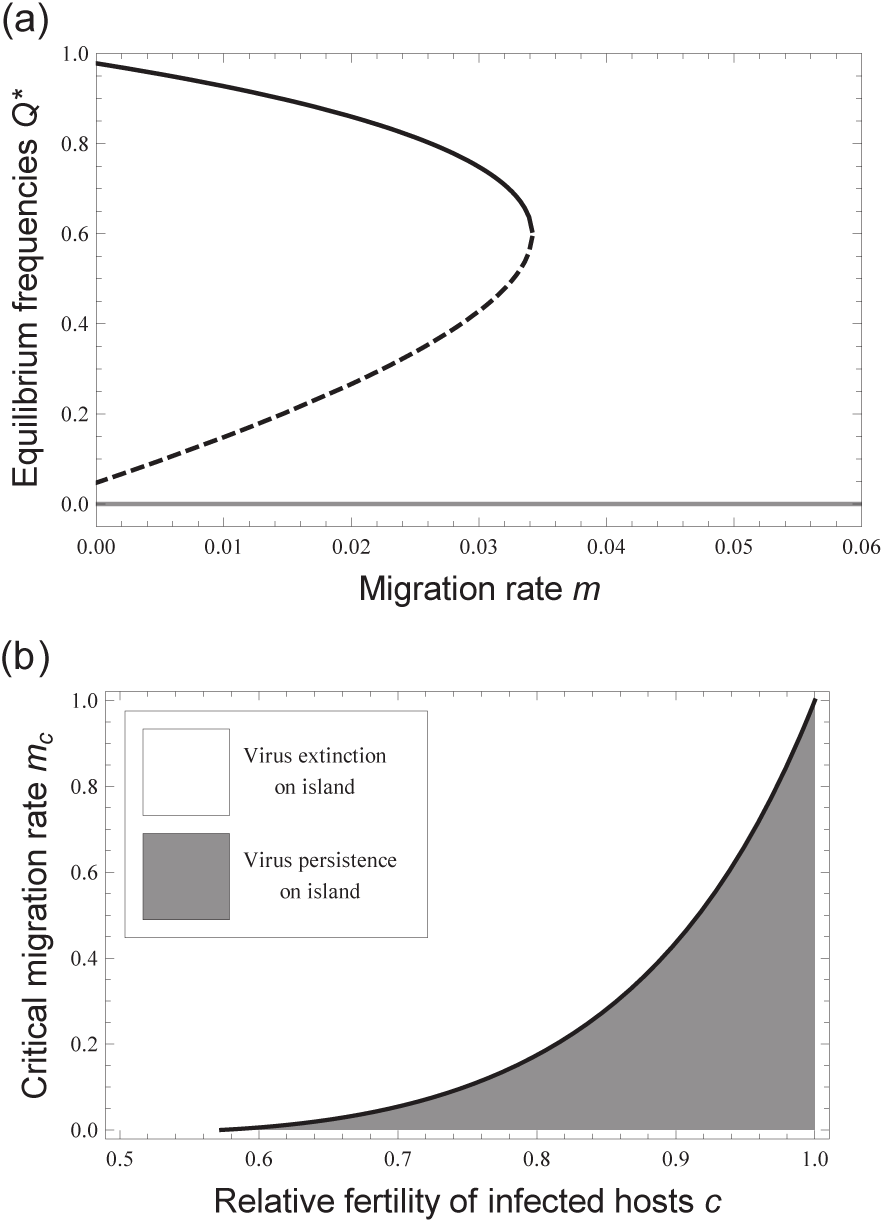
Critical migration rate for the mainland-island model with an un-infected mainland. (a) Virus equilibrium frequencies (*Q*^*^) as a function of the migration rate (*m*) for given parameter 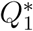 and 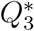, while the dashed line denotes the unstable fixpoint 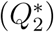. The point where 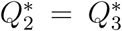 is the critical migration rate, below which the virus may persist in the island population 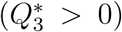, and above which it inevitably becomes extinct on the island 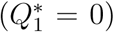. (b) Critical migration rate (*m*_*c*_) as a function of the relative fitness of infected hosts (*c*) for different transmission rates (*t*_*m*_ = 0.99, *t*_*p*_ = 0.495). Below a certain value of *c*, the virus becomes extinct in both populations, whereas above this value a stable infection polymorphism is possible if *m* < *m*_*c*_.

The critical migration rate, *m*_*c*_, is defined as the highest migration rate, below which a stable infection polymorphism is possible. It can be calculated analytically as the migration rate for which 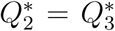. Biologically, the critical migration rate describes the maximum possible immigration for which the virus can persist on the island. For any migration rate above the critical migration rate, there is only a single stable equilibrium frequency, 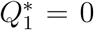, and the virus becomes inevitably extinct in the island population due to migration from the (uninfected) mainland (Fig. 3a). Figure 3b shows the critical migration rate as a function of the relative fitness of infected hosts, *c*. For low values of *c*, it holds that *m*_*c*_ = 0. In general, *m*_*c*_ is a monotonously increasing function of *c*. In summary, the results of this section show that vertically transmitted viruses can persist on an infected island in the face of substantial migration from an uninfected mainland, especially when fertility costs of infection are low.

#### Infected mainland

In this section, the island is assumed to be uninfected and the virus to be at fixation in the mainland population. That means that the viral prevalence in the mainland is 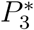, corresponding to the highest stable fixpoint in the single population model. According to the function defined in equation 1, we state the following difference equation describing the viral dynamics in the island population:

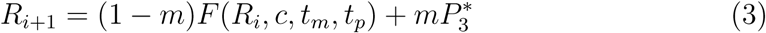

Here, *R*_*i*_ and *R*_*i*+1_ denote the virus frequencies in the island population in subsequent generations, and 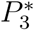 the virus frequency on the mainland. In contrast to the previous sections, we were not able to analyse this model analytically. Therefore, we conducted numerical simulations in order to calculate critical migration rates for different parameter constellations. By varying the migration rate, we determined the critical migration rate (*m*_*c*_) as the highest value below which we observed a stable infection polymorphism. Above the critical migration rate, the simulations showed a viral invasion.

Figure 4 summarizes the results. Three cases can be observed. First, the virus goes to extinction in both populations. This occurs for parameter constellations, for which the virus cannot persist in the single population model. Second, the virus persists on the mainland, and spreads on the island to the same equilibrium frequency. In this case, the only stable equilibrium frequency is positive. Third, the virus persists at high frequency on the mainland, but at low frequency on the island. This case of stable infection polymorphism occurs if parameters are chosen that bistability occurs on the island. Interestingly, it describes a situation, where an infection cannot spread from one population to another despite substantial migration.

**Figure 4:**
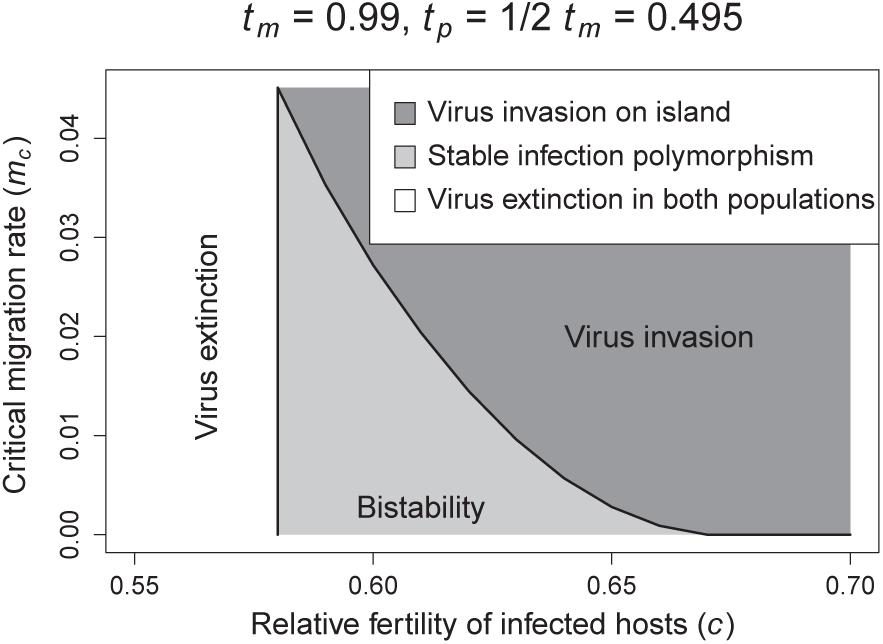
Critical migration rate for the mainland-island model with an infected mainland. Shown is the critical migration rate (*m*_*c*_) as a function of the relative fitness of infected hosts (*c*) for given transmission rates (*t*_*m*_ = 0.99, *t*_*p*_ = 0.495). Values of critical migration rates were calculated by numerical simulations.

A comparison of the two mainland-island models clearly indicates that critical migration rates are higher for the uninfected mainland model than for the infected mainland model. Biologically, this means that an infected island population is more robust to immigration of uninfected individuals than the opposite scenario with an uninfected island and infected migrants.

#### Model with two-way migration

Next, we investigated the infection dynamics for a two-population model with bidirectional migration. A key aspect of the analysis is, under which conditions differences in initial conditions can prevail in the face of migration. As initial conditions we assumed that the virus has spread to its equilibrium in population *A*, but is absent in population *B*. The corresponding migration rates are denoted *m*_*A*_ and *m*_*B*_, respectively.

Let *Q* and *R* denote the virus infection frequencies in population *A* and *B*, respectively. The trans-generational change in virus frequency is described by the following system of two difference equations:

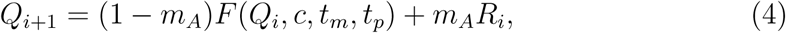

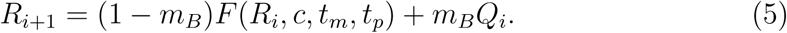

#### Symmetrical migration

We first investigated the infection dynamics when both migration rates are equal (*m*_*A*_ = *m*_*B*_). We screened the parameter space of this model in order to determine under which conditions the virus (1) becomes extinct in both populations, (2) invades both populations, or (3) shows a stable infection polymorphism, where the infection is at a high prevalence in one population and at low prevalence in the other population. We performed numerical estimates for these three zones, by doing spread simulations over 10,000 generations with varying migration rates (Fig. 5). The results show that the first scenario occurs for high and the second for low costs of infection. The third scenario occurs for intermediate costs of infection, and if migration rate is below the critical migration rate.

**Figure 5:**
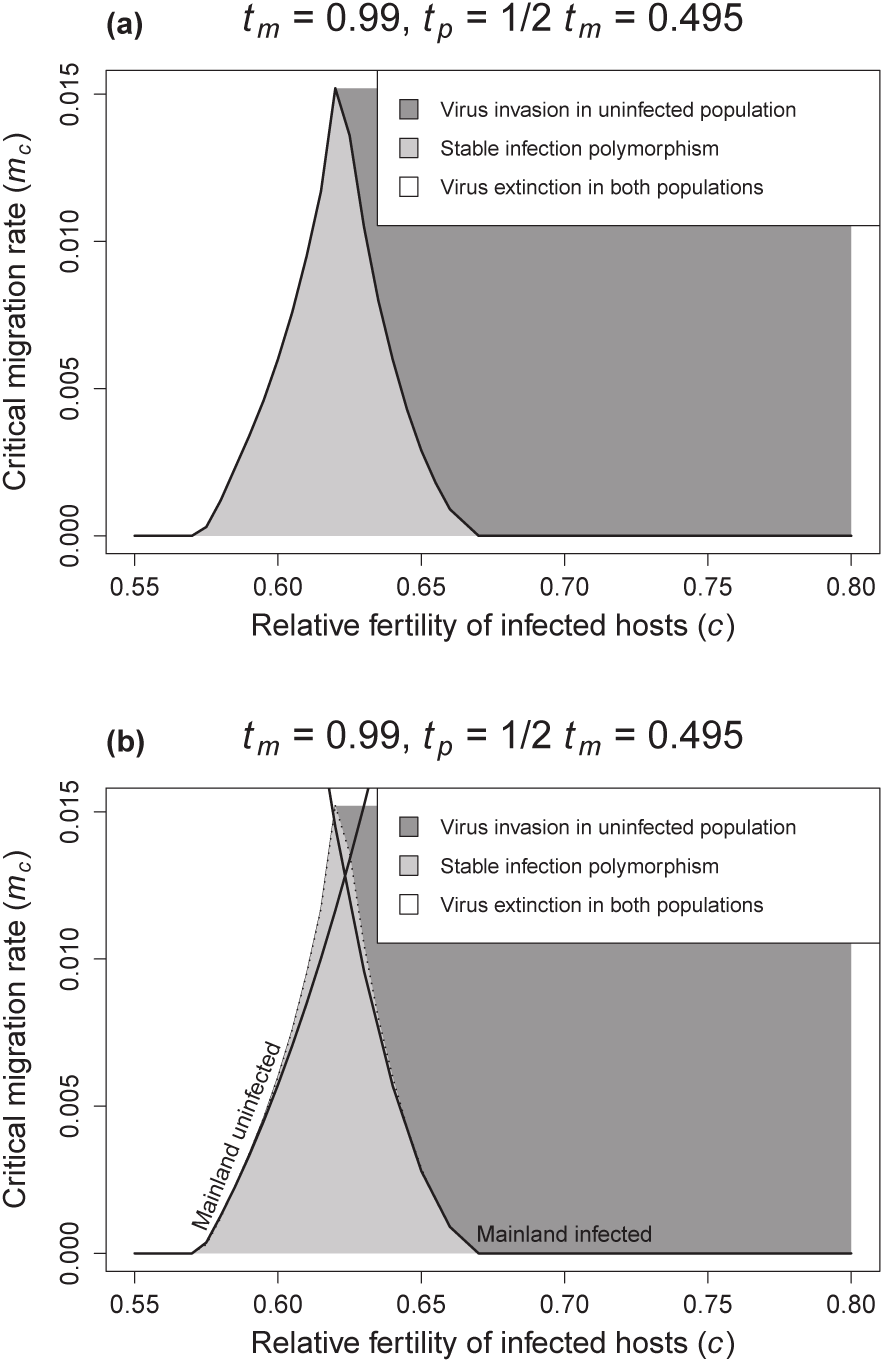
Stability of the infection polymorphism in a two-population model with symmetrical two-way migration. System equilibria were determined by numerical simulations. Both graphs show three regions: (i) a region, where the infection polymorphism is stable (light grey), (ii) a region where the virus spreads and persists in both populations (dark grey), and (iii) a region where the virus goes to extinction in both populations (white). Graph (a) shows numerical simulations. Graph (b) shows in addition the critical migration rates of the corresponding mainland-island models.

Figures 5b shows that the critical migration rates for the mainland-island models are good approximations for the three zones of the two-population model with symmetrical two-way migration. To be more precise, let *m*_*c*,1_ and *m*_*c,2*_ denote the critical migration rates for the two mainland-island models for a certain parameter set (*c, tm, tp*), and *m*_*c*,3_, denote the critical migration rate of the model with bidirectional migration. Then it holds that *m*_*c*,3_ ≤ *Min*{*m*_*c*,1_, *m*_*c*,2_}. Further, *m*_*c*,3_ ≈ *Min*{*m*_*c*,1_, *m*_*c*,2_} is a good approximation for most parameter constellations.

#### Asymmetrical migration

Finally, we analysed the virus infection dynamics when migration rates differ between the populations, and for the initial conditions of population *A* infected and population *B* uninfected. Using computer simulations we found, as above, that the parameter space is separated in three zones: (1) virus extinction in both populations, (2) virus invasion in both populations, and (3) stable infection polymorphism. Figure 6 shows how the zones depend on the migration rates. Interestingly, a stable infection polymorphism is possible for *m*_*A*_ up to 2% and *m*_*B*_ up to 0.3%. This asymmetry reflects the above described effect that the (infected) population *A* is more tolerant to immigration than the (uninfected) population *B*. In other words, immigration results more easily in the spread than in the loss of the virus.

**Figure 6:**
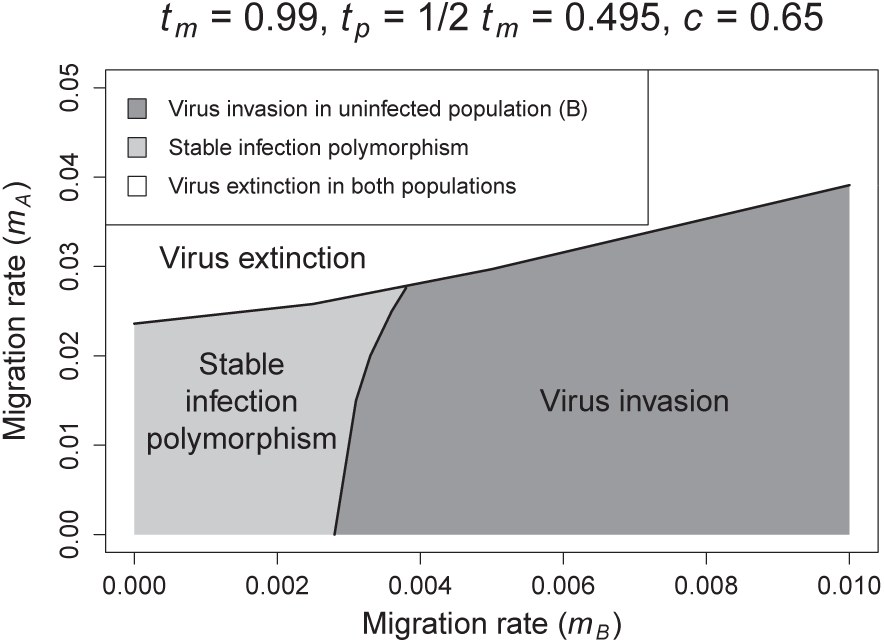
Stability of the infection polymorphism in a two-population model with asymmetrical two-way migration. System equilibria were determined by numerical simulations. The graph shows three regions: (i) a region, where the infection polymorphism is stable (light grey), (ii) a region where the virus spreads and persists in both populations (dark grey), and (iii) a region where the virus goes to extinction in both populations (white).

### 3.4 Discussion

We investigated the infection dynamics of vertically transmitted viruses for different host population structures. Three qualitatively different outcomes occurred in all models: (1) viral spread, (2) no viral spread, and (3) bistability (in the single population model) or stable infection polymorphism (in the two-population models). The key finding of the study is the existence of a zone of bistability for the single population dynamics. Its interesting consequence for the spatial models is that an infection can reach high frequency in one population, but remains at low frequency in a neighbouring population despite substantial migration.

An important question is whether bistability occurs in natural populations. Though our mathematical analysis revealed that bistability occurs only for certain parameter constellations (Fig. 2), empirical evidence for the sigma virus in *Drosophila* suggests that these parameters are biologically realistic. Female and male transmission rates were measured as *t*_*m*_ ≈ 1 and *t*_*p*_ ≈ 0.3 in *D. athabasca* (Williamson, 1961), *t*_*m*_ ≈ 0.92 and *t*_*p*_ ≈ 0.88 in *D. obscura* (Longdon et al., 2011b), and *t*_*m*_ ≈ 0.98 and *t*_*p*_ ≈ 0.45 in *D. affinis* (Longdon et al., 2011b). The fitness reduction of infected hosts is more difficult to measure as sigma virus was shown to have several detrimental effects including fecundity reduction, reduced survival in winter, CO2 sensitivity, and increased hatching time (Williamson, 1961; Seecof, 1964; Fleuriet, 1981). The estimated fecundity reduction of 0.23 in *D. melanogaster* (Seecof, 1964) is therefore an underestimation of the total fitness reduction. Yampolsky et al. (1999) estimated the total fitness reduction of sigma virus in *D melanogaster* as 0.2 *-* 0.3. In summary, a fitness reduction of 0.2 *-* 0.4 seems realistic under natural conditions, and this puts the bistability zone within a realistic range of parameters.

Our analysis of the two population models may help explaining spatial patterns of biparentally inherited parasites. Field surveys of sigma virus revealed large geographical variation in infection prevalence (Williamson, 1961; Wayne et al., 2011; Longdon et al., 2011b). A standard explanation for such patterns is geographical or temporal variation in virus transmission rates or cost of infection, possibly coupled with local extinction and colonization events. This idea is supported by experimental findings that virus transmission rates depend on the nuclear host background and varies geographically (Fleuriet, 1996). Our results show that a stable infection polymorphism is possible for certain parameters, and if migration is below a threshold. This suggests that differences in infection prevalence might happen even if populations do not differ with respect to the parameters describing virus dynamics. We do not want claim that this effect solely explains the described patterns. Rather, we argue that bistability acts in concert with spatial variation of parameters, and that the combination of both creates the geographical patterns of viral prevalence, which are observed in nature.

In our analysis, we have focused on the infection dynamics of biparentally inherited parasites. Another interesting question would be how the presence of infection affects host genome evolution. Though we havent conduced such an analysis, some general conclusions can be drawn from a comparison with uniparentally inherited bacteria *Wolbachia* (Werren et al., 2008). Theoretical analysis suggests that *Wolbachia* infections transform host populations into population genetic sinks if there is immigration from uninfected populations (Engelst ädter and Telschow, 2009; Kobayashi et al., 2011). We expect a similar effect to occur for biparentally inherited viruses if two populations with bidirectional migration are considered, and parameters allow for a stable infection polymorphism. This could result, for example, in a pattern where neighbouring *Drosophila* populations differ with respect to sigma virus frequency, but show no or low divergence at nuclear genes.

In summary, our model analysis shows that biparentally inherited parasites exhibit complex dynamics, including bistability for certain parameter constellations. An important implication is that a stable coexistence between infected and uninfected host populations is possible under biological reasonable conditions, and if migration is below a threshold. This may help understanding geographic patterns of vertically inherited parasites like sigma virus in *Drosophila*.

## 4 Acknowledgments

This work was funded by the Evolutionary Initiative of the Volkswagen-foundation, and the German science foundation (DFG project TE976/2).

## 6 Appendix

### 6.2 A: Graphical illustration of equation 1

**Figure 7:**
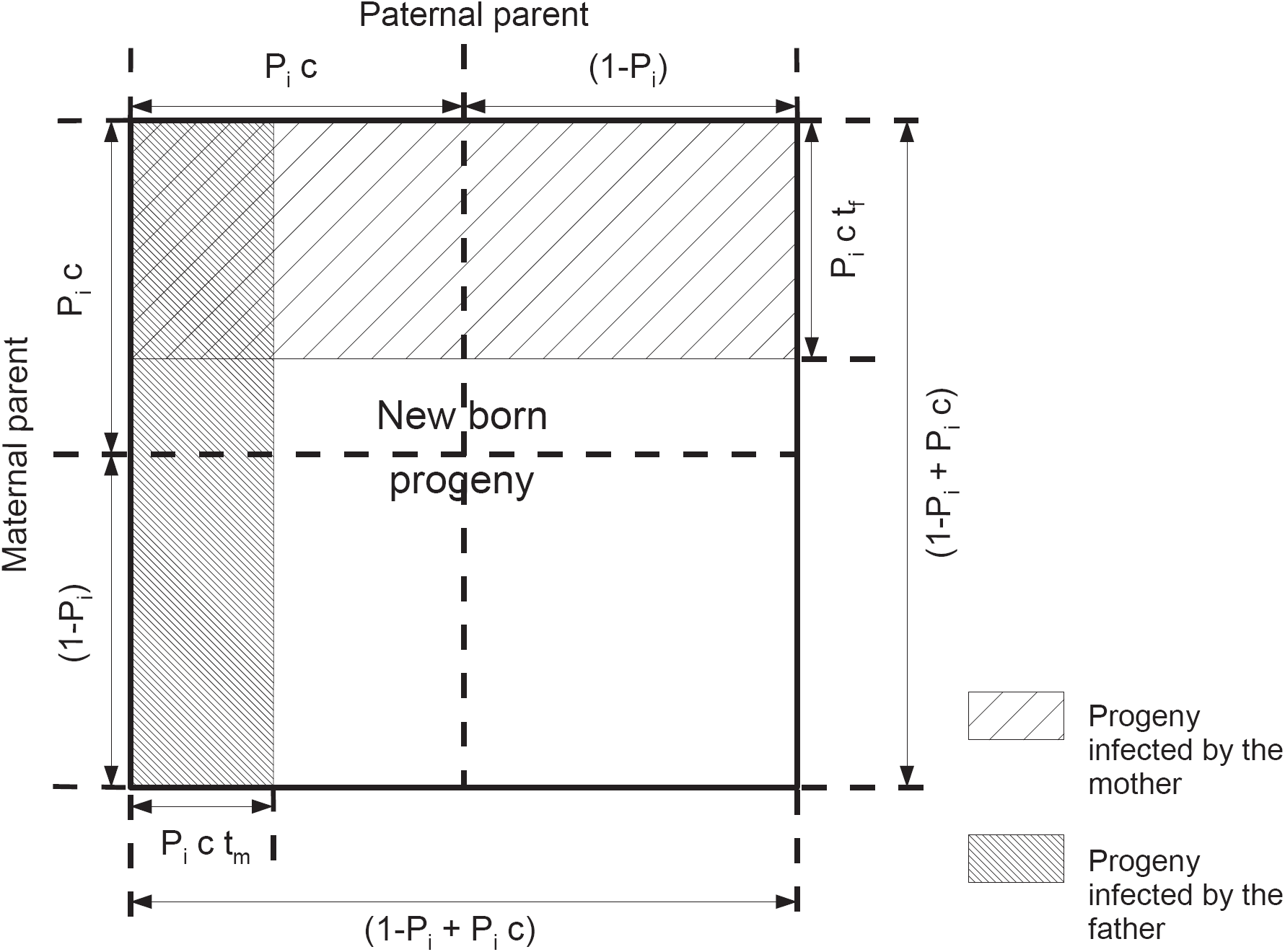
Graphical illustration of equation 1. The Venn diagram square shows the relationship between infection rates of parents and their newborn progeny. *P*_*i*_ is the viral prevalence among both male of female of the parental generation.

### 6.1.1 B: Analytical solutions of fixpoints Single population model

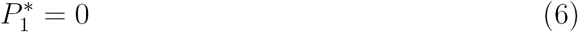

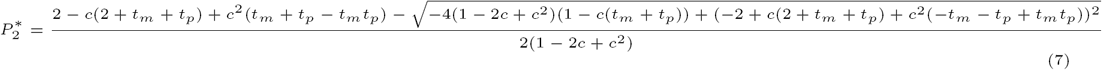

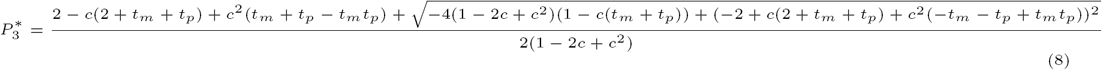

### Mainland island model with uninfected mainland

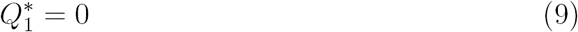

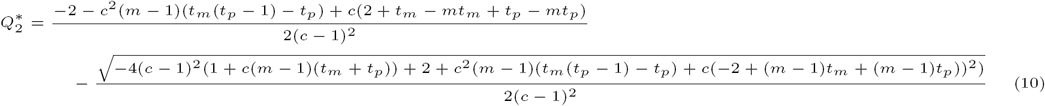

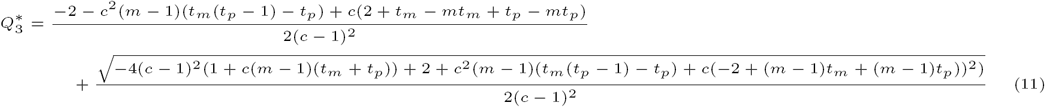

